# Differential effects of α-Synuclein monomers and seeds on the material properties of Tau condensates

**DOI:** 10.64898/2026.04.14.718443

**Authors:** Bineet Sharma, Jinying Wang, Priscilla Chinchilla Retana, Jean Baum, Zheng Shi

**Author notes:** (J.B.); (Z.S.). Campus Straubing for Biotechnology and Sustainability, Technical University of Munich, Schulgasse 16, 94315 Straubing, Germany (Bineet Sharma).

## Abstract

Tau and α-Synuclein (αSyn) frequently co-aggregate in various neurodegenerative disorders. Recently, Tau has been shown to form dynamic, liquid-like condensates that can recruit αSyn, and potentially serve as a precursor to pathological aggregation. However, the quantitative impact of αSyn on the material properties of these condensates remains elusive. Here, we measure the viscosity and interfacial tension of Tau condensates and determine how these properties are modulated by αSyn monomers and fibril seeds. We find that while both forms of αSyn partition efficiently into Tau condensates, they exert vastly different effects on the condensate’s material state. The viscosity of Tau condensates remains unchanged in the presence of αSyn monomers at concentrations up to 200 µM, accompanied by a moderate reduction in the condensates’ interfacial tension. In contrast, the addition of only 5 µM αSyn fibril seeds triggers rapid solidification of Tau condensates, manifested by a nearly 100-fold increase in condensate viscosity within one hour. These findings provide quantitative insights into condensate mechanics, highlighting the unique capacity of αSyn seeds to drive the liquid-to-solid transition of Tau condensates that may underlie the formation of pathological aggregates.

---

Beyond its canonical functions in microtubule assembly & stabilization, axonal transport, and neuronal signaling^1,2^ Tau can misfold and aggregate under pathological conditions, forming paired helical filaments (PHFs) and neurofibrillary tangles (NFTs), a defining histopathological hallmark of Alzheimer’s disease and related tauopathies.^1,3–5^ More recently, Tau has been shown to undergo phase separation under physiologically relevant conditions in vitro and in cells, forming liquid-like condensates in the presence of cofactors, such as RNA, heparin, and crowding agents like polyethylene glycol (PEG).^6–11^ In addition to homotypic assembly, Tau also exhibits heterotypic complex coacervation with proteins such as α-Synuclein (αSyn).^12–16^

The full-length Tau 2N4R (441 residues) is an intrinsically disordered protein containing 2 N-terminal inserts and 4 microtubule-binding repeats that contribute to its multivalent interactions (Figure S1A).^6,7,10^ αSyn is a short 140-residue intrinsically disordered protein composed of an amphipathic N-terminal membrane-binding region, a hydrophobic non-amyloid-β component (NAC) domain that drives aggregation, and an acidic C-terminal tail (Figure S1B). Aggregated αSyn species form amyloid fibrils that constitute a pathological hallmark of synucleinopathies.

In Alzheimer’s disease, αSyn co-pathology has been shown to drive Tau accumulation,^17^ highlighting a functional interplay between these two proteins. The crosstalk between αSyn and Tau has been marked by their colocalization and co-occurrence in human brain tissues.^18–20^ Furthermore, immunohistochemical and biochemical studies involving seeding experiments have revealed that each protein can influence the fibrilization of the other.^21–24^

However, the molecular basis of how αSyn promotes the accumulation and disease transition of Tau remains unclear.

Recent studies have highlighted the role of liquid biomolecular condensates as dynamic, membrane-less entities that can organize cellular biochemical reactions.^25,26^ Liquid condensates formed by aggregation prone proteins, such as Tau, have been shown to undergo maturation to a solidified state, leading to a proposed role of condensates as intermediates in pathological transitions.^27,28^ Recent evidence also suggests that solidified condensates can serve as an amorphous protective state that helps to prevent the formation of ordered fibrillar aggregates.^29–34^ Furthermore, the condensate interface has been shown to play a central role in promoting fibril formation.^31,33,35^

While αSyn is known to partition into Tau condensates,^13,15^ a critical gap remains regarding how different αSyn species such as monomers or fibril seeds (seeds) influence the condensates’ material states. The solidification of liquid condensates into potential pathological states can be quantified as an increase in condensate viscosity, while changes to the condensate interface typically perturb interfacial tension. Therefore, precise measurements of condensate viscosity and interfacial tension are essential to dissect the mechanistic underpinnings of condensate-related pathologies. Recently, we developed the micropipette aspiration (MPA) technique, a direct and label-free method for simultaneous quantifications of the viscosity and interfacial tension of protein condensates in vitro and in cell.^36–39^

In this study, we first quantified the viscosity and interfacial tension of Tau 2N4R condensates using MPA under physiologically relevant conditions. Building on this, we systematically examined the effects of αSyn monomers and seeds on the material properties of Tau condensates. By carrying out these bottom-up analysis, we reveal how αSyn can induce disease-related transitions of Tau, laying the basis for understanding of the crosstalk between synucleinopathies and tauopathies.

We first constructed the phase diagram of purified Tau 2N4R (Figure S1C, S1D) in the presence of PEG-8000 under 25 mM HEPES, 150 mM NaCl, pH 7.4 (Figure 1A, S1D). To minimize experimental variations, we focus on a condition deep in the phase-separated regime (20 µM Tau and 10% PEG-8000) for subsequent condensate material characterizations.

**Figure 1.**
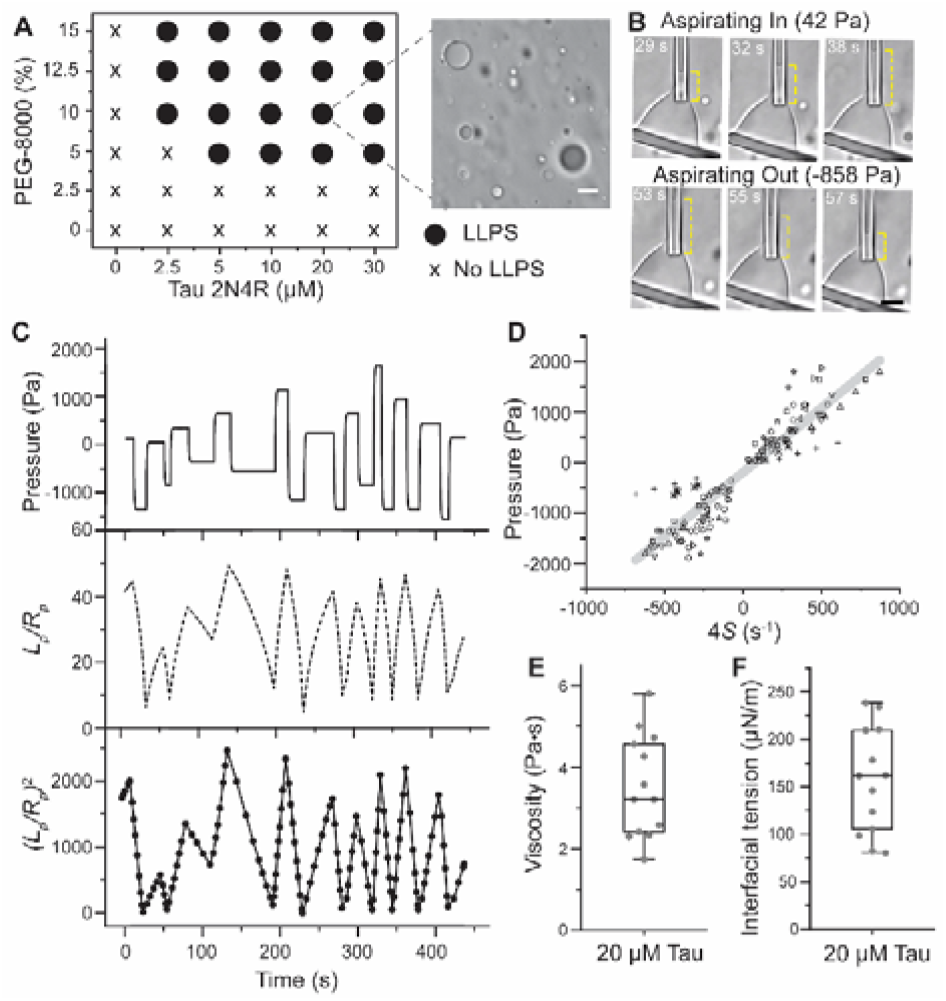
Phase separation of Tau and material properties of Tau condensates. (A) Phase diagram illustrating the formation of Tau condensate with PEG-8000, scale bar 5 µm. (B) Representative transmitted light images showing changes in the aspiration length (*L*_p_) during MPA, scale bar 10 µm. Yellow dash lines trace *L*_p_. (C) Applied aspiration pressure over time (top) and corresponding *L*_p_ normalized by radius of the pipette *R*_p_ (middle). The bottom plot shows the square of *L*_p_/*R*_p_, from which the effective shear rates under each pressure step can be derived from *S* = d (*L*_p_/*R*_p_)^2^/dt. (D) Aspiration pressure vs. 4S for 13 independently prepared condensates. Gray line is a linear fit to all data. (E) Viscosity and (F) interfacial tension of condensates formed by 20 µM Tau in 10% PEG-8000, 25mM HEPES, 150 mM NaCl, pH 7.4. In box plots, the central lines denote the median, the boxes span the interquartile range (25 to 75%), and the whiskers extend to 1.5 times the interquartile range, same below.

Next, we quantified the material properties of Tau condensates using MPA. Under the selected condition (20 µM Tau, 10% PEG-8000), Tau condensates behave as a Newtonian liquid (Figure 1B, 1C), with a well-defined effective shear rate *S* that changes linearly with the aspiration pressure *P*_asp_ (Figure 1D). Following eq (1), the slope and intercept of *P*_asp_ vs. *S* give the viscosity and interfacial tension of the aspirated condensate, respectively. The viscosity of Tau condensates was 3.5 ± 1.2 Pa⍰s (mean± s.d.; same below) and the interfacial tension was 156 ± 55 µN/m (Figure 1E, 1F). We noticed that impurities in recombinant Tau samples can reduce the condensate’s viscosity and interfacial tension (Figure S1E, 1F). In this study, we used the highest purity fractions of Tau obtained from each purification.

We next examined the effect of αSyn on pre-formed Tau condensates. Confocal fluorescence microscopy was used to visualize the partitioning of labelled Tau-AF488 and αSyn-AF647 into condensates. Enrichment was quantified by the partition coefficient (PC), defined as the ratio of the protein fluorescence in the dense phase to that in the dilute phase. Fluorescent labeling was only used for partitioning measurements and all MPA experiments were performed with unlabeled proteins to avoid photo-artifacts.

Monomeric αSyn partitioned strongly into Tau condensates, as reflected by the high PC values (Figure 2A, Figure S2A). PC decreased from ∼120 to ∼45 as the total concentration of αSyn monomers increased from 5 to 200 µM, consistent with simple, non-cooperative recruitment of αSyn into Tau condensates. Notably, under these buffer conditions, 200 µM αSyn monomers alone do not form condensates.^40^

**Figure 2.**
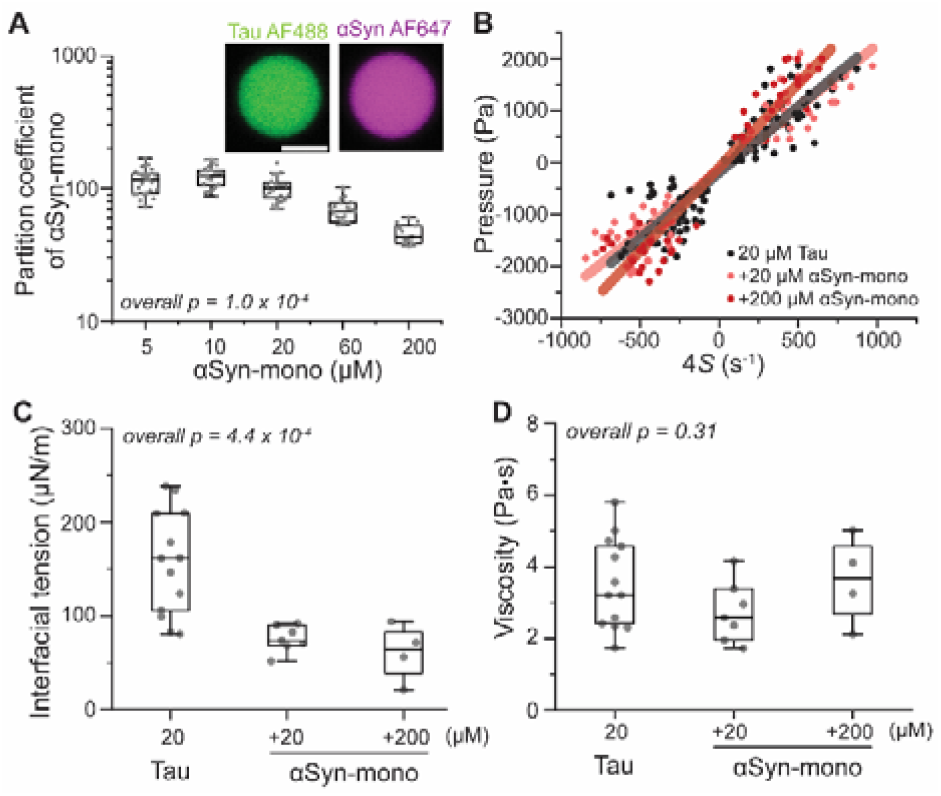
Effect of αSyn monomers on the material properties of Tau condensates. (A) Partition coefficient of αSyn monomers into Tau condensates where Tau is labelled with Tau-AF488, and αSyn with αSyn-AF647. Inset shows representative fluorescence images of the condensate, scale bar 2 µm. (B) Pressure vs. 4S plots of Tau condensates in the presence of αSyn monomer as measured by MPA. (C) Change in interfacial tension and viscosity (D) of Tau condensates with increasing αSyn monomer concentration. *p* values were determined using one-way analysis of variance (ANOVA).

We estimate that the concentration of αSyn monomers in the dense phase is ∼5mM (Figure S2B, S2C), which could have a prominent impact on the rheology of Tau condensates. To test this, we performed MPA measurements on Tau condensates supplemented with αSyn monomers. While the interfacial tension of Tau condensates decreased 3-fold upon the addition of monomeric αSyn (Figure 2C), the viscosity of Tau condensates remained surprisingly stable (Figure 2B, 2D), despite the presence of millimolar level of αSyn monomers in the condensates. Interestingly, the strong partitioning of αSyn monomers also did not significantly affect the concentration of Tau in the dense phase (Figure S2D, S2E).

We next evaluated the effect of αSyn seeds on Tau condensates. In contrast to the intrinsically disordered monomer, αSyn seeds derived from preformed fibrils adopt β- sheet rich cores with flanking disordered termini.^41–43^ The αSyn seeds used here have an average length of 50 nm and average height of 5 nm (Figure S3A-3C). Like monomers, αSyn seeds strongly partitioned into Tau condensates, with PC ∼100 (Figure 3A, 3B). Notably, seeds showed heterogeneous accumulation in condensates at 5 µM monomer equivalent concentration (Figure 3B). In contrast to monomers, αSyn seeds substantially increased the viscosity of Tau condensates (Figure 3C, 3D). Even at 1 µM concentration, seeds increased the viscosity of Tau condensates by ∼5-fold. Increasing the seeds concentrations to 5⍰µM resulted in a ∼30-fold increase in condensate viscosity on average, while the interfacial tension of condensates was not measurably affected. Importantly, these effects of seeds occurred at αSyn concentrations (≤5 µM) far lower than those used for monomers (up to 200 µM).

**Figure 3.**
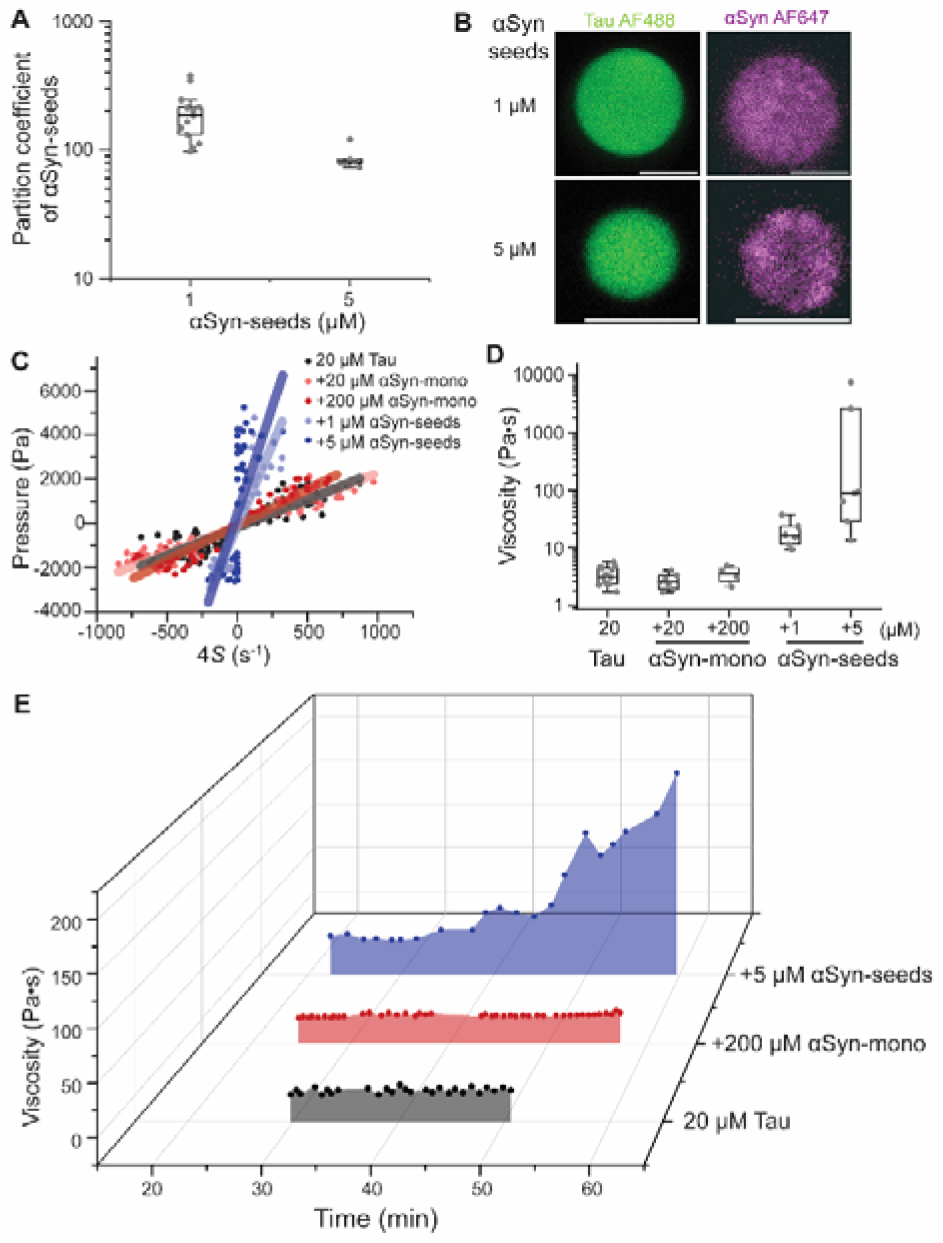
Aging effect of αSyn-seeds on Tau condensates. (A) Partitioning of αSyn seeds into Tau condensates and (B) representative images of Tau condensate supplemented with αSyn seeds where Tau is labelled with Tau-AF488, and seeds were made from αSyn monomers containing αSyn-AF647, scale bar 5 µm. (C) Pressure vs. 4S plot of Tau condensates in the presence of αSyn monomers or seeds. (D) Viscosity of Tau condensates with increasing concentration of αSyn monomers and seeds. (E) 3D plot showing the time-dependence of Tau condensate viscosity without αSyn (black), with 200 µM αSyn monomers (red), or with 5 µM αSyn seeds (blue).

Tracking the viscosity of Tau condensates over one hour, we observed that αSyn seeds, but not monomers, led to a continuous increase in the measured viscosity (Figure 3E), corresponding to an apparent aging of condensates. To determine whether this effect was associated with fibril formation in condensates, we performed a Thioflavin T (ThT) assay on Tau condensates supplemented with αSyn seeds. Consistent with previous studies, we observed a basal level of ThT fluorescence in Tau condensates that did not change with the addition of αSyn seeds.^44^ More importantly, the fluorescence of ThT did not measurably increase after incubating Tau condensates with αSyn seeds for 2 hours (Figure S3D). These results suggest that the increase in the viscosity of Tau condensates (within 1 hour of adding αSyn seeds) could not be attributed to fibril formation within the condensates.

Tau and αSyn are known to interact by electrostatic interactions where positively charged proline-rich domains of Tau interact with the negatively charged C-terminal region of αSyn.^13,15,16^ At physiological pH (7.4), Tau carries a net positive charge (+2) while αSyn is substantially anionic (−9 charge), which provides a clear basis for heterotypic association. The decrease in the interfacial tension of Tau condensates under high concentrations of αSyn (Figure 2C) is consistent with established charge-driven alterations of the interfacial properties in heterotypic coacervates.^45,46^

Despite efficient partitioning, monomeric αSyn does not change the concentration of Tau in the dense phase (Figure S2), indicating that αSyn behaves as a client while Tau serves as the primary scaffold for the condensates.^13^ Moreover, the observation that αSyn monomers do not affect the viscosity of Tau condensates (Figure 2D) suggests that αSyn monomers interact weakly and do not disrupt the existing internal network formed by the Tau scaffold.^47^ Within the framework of dominance analysis,^50^ αSyn monomers can be classified as a non-dominant client that does not impact the internal network interactions.

In contrast to its monomeric form, αSyn seeds substantially increase the viscosity of Tau condensates (Figure 3). Fibrillar αSyn contains a structured amyloid core spanning approximately residues L38–K97,^41–43^ flanked by disordered N- and C-termini that form a dynamic “fuzzy coat”.^51^ While the core provides structural stability, the fuzzy coat mediates key functions^52,53^ including monomer recruitment,^49,54,55^ secondary nucleation processes,^56^ fibril cellular transmission,^57^ interactions with other proteins,^58–60^ and serves as a potential therapeutic target.^61^ In particular, the highly negatively charged C-terminus remains solvent-exposed^41,62^ along the fibril surface, creating a multivalent binding platform for positively charged biomolecules such as Tau. As recently demonstrated, this multivalency allows a single seed to simultaneously engage multiple Tau molecules with a higher overall avidity compared to the individual affinity of αSyn monomers.^47^ The multivalent interactions between αSyn seeds and Tau effectively strengthens the condensate scaffold by providing additional interactions, which is directly reflected in the observed elevation in condensate viscosity.^63–65^

Together, our findings demonstrate how distinct αSyn species differentially reprogram the material properties of Tau condensates. Particularly, αSyn seeds lead to a rapid maturation of Tau condensates, a transition beyond the capacity of αSyn monomers. These observations begin to address a critical gap in our understanding of neurodegeneration by providing a physicochemical framework for how Tau–αSyn crosstalk in neurodegenerative diseases may occur through modulations of condensate material properties.

## Displayed equations

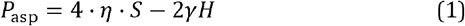

Where *P*_*asp*_ is the aspiration pressure, *η* is the viscosity of the aspirated condensate, *γ* is the interfacial tension of the condensate, *H* is the radius of curvature of the condensate interface inside pipette. *S* is the effective shear rate defined as S = *d* (*L*_p_/*R*_p_)^2^/dt, where *L*_p_ is the length of the condensate aspirated inside the micropipette and *R*_p_ is the radius of the micropipette. The equation assumes that the characteristic size of the condensate outside the pipette is much larger than *R*_p_.

## ASSOCIATED CONTENT

## AUTHOR INFORMATION

### Author Contributions

**Conceptualization**: Z.S., J.B.

**Methodology:** B.S., Z.S.

**Investigation:** B.S. (all the experiments including MPA experiments; Tau expression, purification, and labeling of Tau and αSyn, partitioning experiments), J.W. (Tau MPA measurements, ThT assays), P.C.R. (αSyn monomer purification, seed preparation, AFM imaging).

**Formal analysis:** B.S., Z.S.

**Project administration**: Z.S., J.B.

**Funding acquisition**: Z.S., J.B.

**Supervision:** Z.S.

**Writing –** original draft: B.S.

**Writing** – review & editing: B.S., Z.S., with input from all authors. All authors have approved the final version of the manuscript.

### Funding Sources

We thank funding supported by the National Institute of General Medical Sciences of the National Institutes of Health grant R35GM147027 (Z.S.) and R35GM136431 (J.B.). P.C.R. acknowledges funding from the National Institutes of Health under the T32 Training in Translating Neuroscience to Therapies grant T32NS115700.

## ACKNOWLEDGMENT

We thank Jerelle Joseph, Yashraj Wani, and Nate Hess for helpful discussions.

## Notes

### Competing Interest Statement

The authors have declared no competing interest.

