## Supporting Information for "Differential effects of α-Synuclein monomers and seeds on the material properties of Tau condensates"

The supporting information includes:

Methods and Materials

Supplementary Figure captions

Supplementary Figure S1 to S3

**Methods and Materials**

**Recombinant Tau 2N4R expression and purification**

The plasmid pET-29b(+) encoding full-length wild-type human Tau (2N4R) was transformed into *Escherichia coli* BL21(DE3) strain and was plated on Luria Broth (LB) agar supplemented with 50 µg/mL kanamycin and incubated overnight at 37°C. A single colony was then inoculated into a starter culture with 50 mL of LB broth containing 50 µg/mL kanamycin and grown overnight at 37°C with shaking at 180 rpm (Scientific Innova 44 Incubator Shaker, New Brunswick, NJ). 1-2 volume % of the starter culture was then scaled up to 2 L culture and grown again at 37°C at 180 rpm in the presence of kanamycin until OD at 600 nm reaches between 0.5-0.6. For Tau 2N4R expression, *E. coli* culture was induced by 0.5 mM IPTG for 2 hours at 37 °C. Cells with over expressing Tau were collected at 11,500 × g for 15min at 4 °C. The pellet was resuspended in lysis buffer containing 20 mM MES pH 6.8, NaCl 500 mM, EDTA 1 mM, MgCl_2_ 0.2 mM, DTT 5 mM with protease inhibitor cocktail tablet. A homogeneous pellet in lysis buffer was then sonicated on ice at 30% amplitude with 10 s on and 20 s off for 5 minutes (Sonicator Dismembrator Model 500, Fisher Scientific, MA). Afterwards, the cell lysate was heated over boiling water for 25 minutes and then cooled on ice and centrifuged at 125,000 × g at 4 °C for 50 min using Optima MAX-TL ultracentrifuge. The supernatant was dialyzed with 2 L dialysis buffer (MES 20 mM pH 6.8, NaCl 50 mM, EDTA 1 mM, MgCl_2_ 2 mM, DTT 2 mM) in a 3.5 kDa snakeskin (Thermo) overnight (~16-18 hours). Tau 2N4R was purified in two steps, one with 2 × 5 mL cationic exchange column (HiTrap SPFF, Cytiva, MA, USA) and other with size exclusion chromatography. Cation exchange column was equilibrated with equilibration buffer (MES 20 mM pH 6.8, NaCl 50 mM, EDTA 1mM, MgCl_2_ 2 mM, DTT 2 mM before sample injection (dialyzed). Tau containing lysate after dialysis was filtered through a 0.22 μm PVDF filter and injected into the column at a flow rate of 1 mL/min. Tau protein was eluted gradually between 0 – 60% of elution buffer (MES 20 mM pH 6.8, NaCl 1 M, EDTA 1 mM, MgCl_2_ 2 mM, DTT 2 mM). The fractions containing expected MW tau (confirmed by SDS-PAGE) were pooled and injected again into a size exclusion column (Superdex 75 10 300 GL) with a flow rate of 0.5 mL/min using equilibration buffer. Fractions containing tau analyzed by SDS-PAGE were then buffer exchanged to a desired buffer and stored at -80 °C. Tau concentration was measured using NanoDrop using MW 45849 and extinction coefficient of 7575 M^-1^cm^-1^.

**Recombinant α-Synuclein expression and purification**

Expression of non-acetylated human αSyn occurred via transformation into *E. coli* BL21(DE3) cells (New England Biolabs, MA) and plated onto agar plates containing the antibiotic ampicillin (50 µg/mL). 50 mL starter cultures were inoculated using one colony each and grown in LB media at 37°C, 160 rpm overnight (Scientific Innova 44 Incubator Shaker, New Brunswick, NJ). 1L LB cultures were inoculated using 10 mL starter cultures and were allowed to grow at 37°C, 180 rpm until an O.D of 0.6 was reached. At this point, the cultures were induced with 1 mM IPTG and grown at 20°C, 180 rpm for 16-18 hours. After overexpression, cultures were pelleted down and stored at -80 °C until use.

Non-acetylated human αSyn was purified by resuspending the bacterial pellets in 10 mM PBS (pH 7.4). The protein solutions were lysed via sonication (Sonicator Dismembrator Model 500, Fisher Scientific, MA) at 30% amplitude for 8 cycles of 15 seconds on 30 seconds off. Solutions were kept on ice to prevent degradation. Following lysis, samples were boiled for 20 minutes and centrifuged at 20,000 rpm for 45 minutes (Avanti J-26S XPI Centrifuge, Beckman Coulter, CA) to remove cellular debris. The supernatants were collected and treated with streptomycin sulfate (10 mg/mL) to remove nucleic acids. The solutions were mixed at 4°C for 20 minutes and centrifuged as previously described. The protein was precipitated by add-ing ammonium sulfate (0.361 g/mL) and mixed at 4°C for 1 hour after which the solutions were centri-fuged once more to collect the final protein pellets. The pellets were resuspended in Buffer A (32 mM Tris, pH 7.8) and prepared for fast protein liquid chromatography (FPLC) by filtration using a 0.22 µm syringe filter. The filtered protein solutions were injected onto a HiTrap Q HP anion exchange column (Cytiva AKTA, MA) and the purified protein was eluted at 50% Buffer B (32 mM Tris, 500 mM NaCl, pH 7.8) using an AKTA Pure FPLC (Cytiva AKTA, MA). The purified protein was flash frozen using LN2 and stored at -80 °C until use.

**α-Synuclein** **fibril** **seeds preparation**

Flash-frozen αSyn was thawed on ice, and large aggregates were removed using a 50 kDa centrifugal filter (Millipore Sigma, St. Louis, MO). The protein was then concentrated into 10 mM PBS (pH 7.4) using a 3 kDa centrifugal filter (Millipore Sigma) to a final concentration of 300–500 µM. 500 µL of the concentrated protein was aliquoted into a 1.5 mL microcentrifuge tube and incubated in an orbital thermomixer (Thermomixer C, Eppendorf, MA) at 37 °C with shaking at 1000 rpm for 7 days. Fibrils were collected by centrifugation at 15,000 rpm for 2 hours (Microcentrifuge 5430, Eppendorf, MA). The resulting pellets were washed via multiple rounds of resuspension in 10 mM PBS (pH 7.4) followed by centrifugation at 15,000 rpm for 2 hours to remove residual soluble and non-fibrillar species. Fibril pellets were stored at room temperature until use. Fibril seeds were prepared by sonicating 1 mg/mL of fibrillar protein (15 seconds on, 15 seconds off, 30% amplitude) for 3 cycles on ice. AFM samples were prepared by diluting seeds at a 1:10 concentration. The concentrations of fibrils and seeds were determined by dissociating in guanidinium and performing a bicinchoninic acid assay. The concentration is then reported in monomer equivalents.

**Air AFM imaging**

For the air AFM imaging sample preparation, 50 µL of fibril seeds were placed onto freshly cleaved mica and allowed to bind to the surface for approximately 5 minutes. After, the mica surface was washed with 2 mL of Milli-Q water, and the sample was dried at room temperature for 2 hours in a laminar flow hood. After drying, the sample was placed into the AFM for imaging (Cypher ES AFM, Asylum Re-search, Oxford Instruments, United Kingdom). Imaging was performed at 25°C with AC240 tips (nominal resonance frequency of 70 kHz, nominal spring constant of 2 N/m).

**Tau phase separation in presence of a crowding agent**

The phase separation of Tau was induced by supplementing polyethylene glycol (PEG-8000, Thermo Fisher Scientific) as a crowding agent in HEPES buffer (25 mM HEPES, 150 mM NaCl, pH 7.4) at room temperature. As shown in Figure 1, Tau with varying final concentration of PEG-8000 were mixed in a low binding tubes by slow pipetting and observed under microscope using homemade observation chamber fabricated by sandwiching two cover glasses with a layer of vacuum grease in between. Tau condensates were observed using Nikon Ti2-A inverted fluorescence microscope with 60 x objective unless otherwise mentioned.

**Fluorescent labelling of Tau and** **α-Synuclein**

The presence of two native cysteines in Tau at 291 and 322 were site-specifically labeled by maleimide-thiol reaction strategy using AlexaFluor488 C5-maleimide following manufacturer’s protocol (Thermo Fisher Scientific). Briefly, purified Tau was buffer exchanged into labelling buffer (25mM HEPES, 150mM NaCl, pH 7.4) using Zeba™ Spin Desalting columns (Thermo Fisher Scientific). Tau and AF488 were mixed in the molar ratio of 1:1000 and incubated overnight at 4 °C. The excess dye was removed using Zeba™ spin columns.

Similarly, for αSyn, single cysteine mutation at A140C was employed for site-specific labeling with AlexaFluor647 C2-maleimide in the molar ratio of 1:1000 and incubated overnight at 4 °C. The excess dye was removed by Zeba™ spin columns and concentration of labelled proteins were measured using NanoDrop with absorbance correction based on manufacturer’s protocol.

**Partitioning of α-Synuclein** **monomers & seeds** **into Tau condensates**

Tau condensates were freshly prepared by mixing 20 µM Tau (15 µM unlabeled Tau: 5 µM labeled Tau-AF488) with 10% PEG-8000 in HEPES buffer and then supplemented with varying concentration of αSyn monomers (x µM unlabeled αSyn: 5 µM labeled αSyn-A140C-AF647). The protein mixed solution was transferred to a glass slide precoated with 5% Pluronic acid (F-127, Sigma Aldrich). An observation chamber was created by employing a silicone spacer of 0.5 mm and condensates were observed under confocal microscopy, Zeiss Axio Observer 7 inverted microscope equipped with an LSM900 laser scanning module. Similarly, partitioning of labelled preformed α-Synuclein seeds was conducted under identical conditions. All images were analyzed using Fiji ImageJ.

**Thioflavin T (ThT) fluorescence assay**

A stock solution of Thioflavin T (ThT; Thermo Scientific Chemicals, CAS 2390-54-7) was prepared in HEPES buffer (25 mM HEPES, 150 mM NaCl, pH 7.4). For this assay, unlabeled preformed αSyn fibril seeds, Tau, and ThT were combined in desired concentrations in the presence or the absence of PEG-8000 (Figure S3D). ThT was added last to initiate the assay with a final concentration of 20 µM in all samples. ThT assay was performed under four conditions: (i) αSyn seeds + ThT, (ii) Tau + αSyn seeds + ThT, (iii) Tau + PEG-8000 + ThT, and (iv) Tau + αSyn seeds + PEG-8000 + ThT. Samples were imaged using Nikon Ti2-A inverted fluorescence microscope at room temperature for 2 h.

**Micropipette aspiration (MPA)**

In this method, micropipettes were fabricated using a thin-walled glass capillaries TW100-4 (OD = 1.0 mm, ID = 0.7 mm, length = 100 mm) by a precision puller instrument (PUL-1000, World Precision Instruments). Each capillary tube yielded two fine capillaries. The tip of the capillary was then carefully trimmed to form an orifice, typically ranging between 2 to 5 µm in diameter. Following this, the tip was bent to an angle of ~ 60° using a microforge device (DMF1000, World Precision Instruments) to help in positioning during MPA experiments. This finely tuned micropipette was then filled with appropriate buffer solution and attached with external pressure pump (FLUIGENT, ESORT-PCK01). Before each aspiration experiment, zero pressure was carefully measured using milli-Q water in a glass-bottom dish.

MPA experiments were performed on Nikon Ti2-A inverted fluorescence microscope with 60 x objective equipped with motorized stage and two micromanipulators (PatchPro-5000, Scientifica), an optical tweezers (Tweez305, Aresis) and a pressure pump (FLUIGENT, ESORT-PCK01). In the experimental setup, protein condensates were transferred to a glass bottom dish mounted over the microscope. One pipette with fused orifice acted as a holding pipette to immobilize the protein condensate for better aspiration recording, while another pipette was connected to pressure flow control set up. Tau condensates were fused together with the help of optical tweezers to achieve a condensate size of >10 µm in diameter, typically 3-5 times larger than orifice of the capillary. The MPA data was collected using positive and negative pressure profiles regulated by external pressure pump and analyzed for biophysical properties. To minimize sample evaporation, a local humidity environment was created by adding Milli-Q water to the outer region of the glass-bottom dish.

Several preformed micron-size Tau condensates were merged using optical tweezers to generate a condensate of ≥ 10 µm in diameter (see method section). This single condensate was positioned on a sealed holding capillary and aspirated into another ~3 µm glass micropipette under external pressure. Positive pressure was applied to aspirate the condensate into the capillary, called “aspirating-in” while negative pressure pulled it back called “aspirating-out”, generating a corresponding change in the aspirated length (Figure 1B).

**Supplementary Figure captions**

**Supplementary Figure 1.** **Domain architecture, charge distribution of αSyn & Tau, and Tau 2N4R phase diagram.** (A) Schematic representation of Tau 2N4R domains, including N-terminal inserts (N1, N2), proline-rich regions (P1, P2), and microtubule-binding repeat domains (R1–R4). The net charge per residue (NCPR) profile highlights alternating positively (blue) and negatively (red) charged regions along the sequence. (B) Domain organization of αSyn showing the amphipathic N-terminal region, hydrophobic NAC domain, and acidic C-terminus, with the corresponding NCPR profile indicating charge segregation. (C) SDS-PAGE analysis of Tau 2N4R fractions, showing purified (“pure”) and less purified (“impure”) samples. (D) Bright-field microscopy images showing phase separation of Tau 2N4R as a function of protein concentration (0–30 µM) and PEG-8000 concentration (0–15%). Highlighted image represents 20 µM Tau with 10% PEG-8000 used in this study, scale bar, 10 µm. (E) Comparison of bulk viscosity measurements of Tau pure fraction to the impure fraction (*p* = 1.8 × 10⁻⁴). (F) Interfacial tension measurements reveal reduced surface tension in the impure fraction relative to the pure fraction (*p* = 0.009), indicating altered material properties.

**Supplementary Figure 2. αSyn partitions into Tau condensates in a concentration-dependent manner.**

(A) Confocal fluorescence images of mixed condensates formed by 20 µM Tau (15 µM unlabeled and 5μM AF488-labeled) and increasing concentrations of αSyn (5–200 µM, with fixed 5 µM AF647-labeled αSyn). αSyn efficiently enriches within Tau droplets with increasing concentration, while Tau remains robustly partitioned across conditions. Scale bars, 2 µm. (B, C) Quantification of effective fluorescence intensity of αSyn in the dense (B) and dilute (C) phases as a function of αSyn concentration, showing enhanced partitioning into condensates at higher concentrations. (D, E) Corresponding quantification of Tau fluorescence intensity in the dense (D) and dilute (E) phases. Tau enrichment in the dense phase and depletion from the dilute phase remain relatively stable across increasing αSyn concentrations, indicating minimal perturbation of Tau partitioning. From Figure (B) & (C), fluorescence intensities were converted to concentrations using a linear calibration factor, assuming saturation exists at 200 µmM αSyn, (*C* = *I*_dense/dil_ /*I*_200_ × 200 µmM), yielding an estimated ~5 mM enrichment of αSyn in the dense phase. Where *C* = estimated αSyn concentration, *I*_dense/dil_ = average fluorescence intensity of dense or dilute phase, *I*_200_ = average fluorescence intensity at 200 µmM αSyn.

**Supplementary Figure 3.** **Thioflavin T (ThT) assay of Tau under PEG-induced phase separation in the presence of αSyn seeds.**

(A) Atomic force microscopy images of αSyn fibril seeds, with an average length of 49.2 ± 23.6 nm (B) and height of 4.74 ± 0.72 nm (C) (mean ± s.d.), scale bar 400 nm. (D) Fluorescence images of ThT assays of Tau (20 µM) in the absence and presence of αSyn seeds (5 µM) and PEG-8000 (10%), recorded at *t* = 0 and 120 min. Under PEG-induced phase separation, Tau condensates exhibit a baseline ThT signal that does not show a significant increase upon addition of αSyn seeds, indicating no substantial enhancement of ThT-positive species within condensates.

**Figure S1**

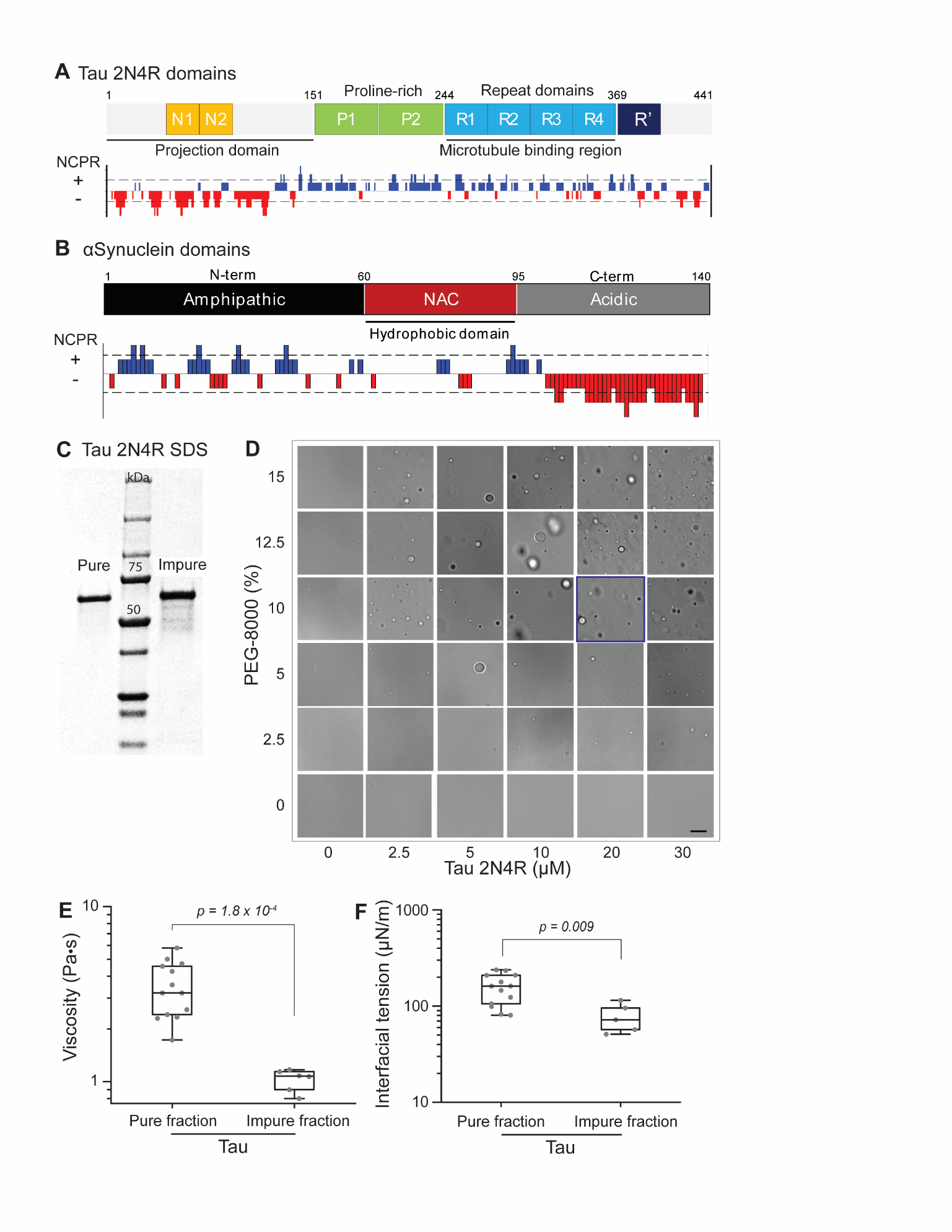

**Figure S2**
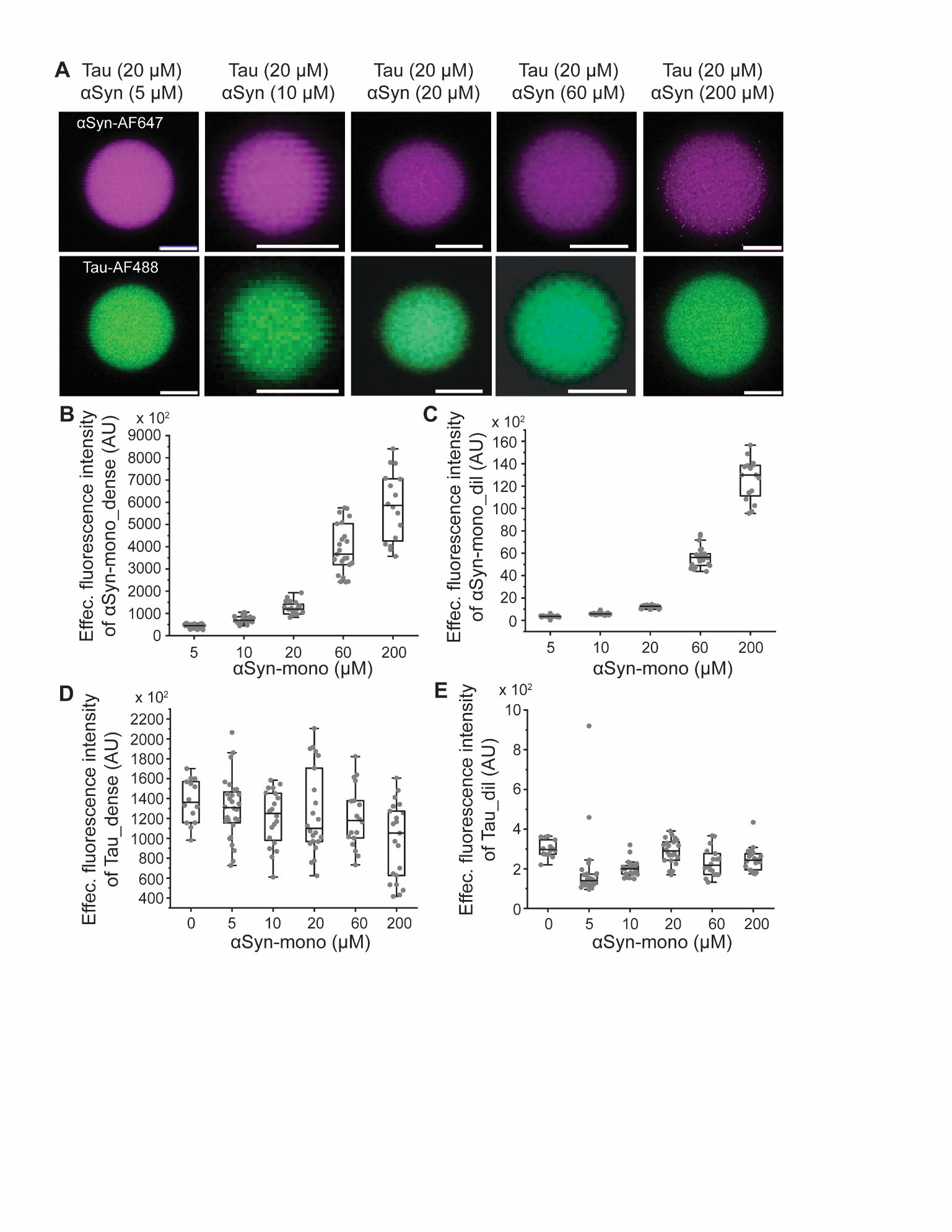

**Figure S3**

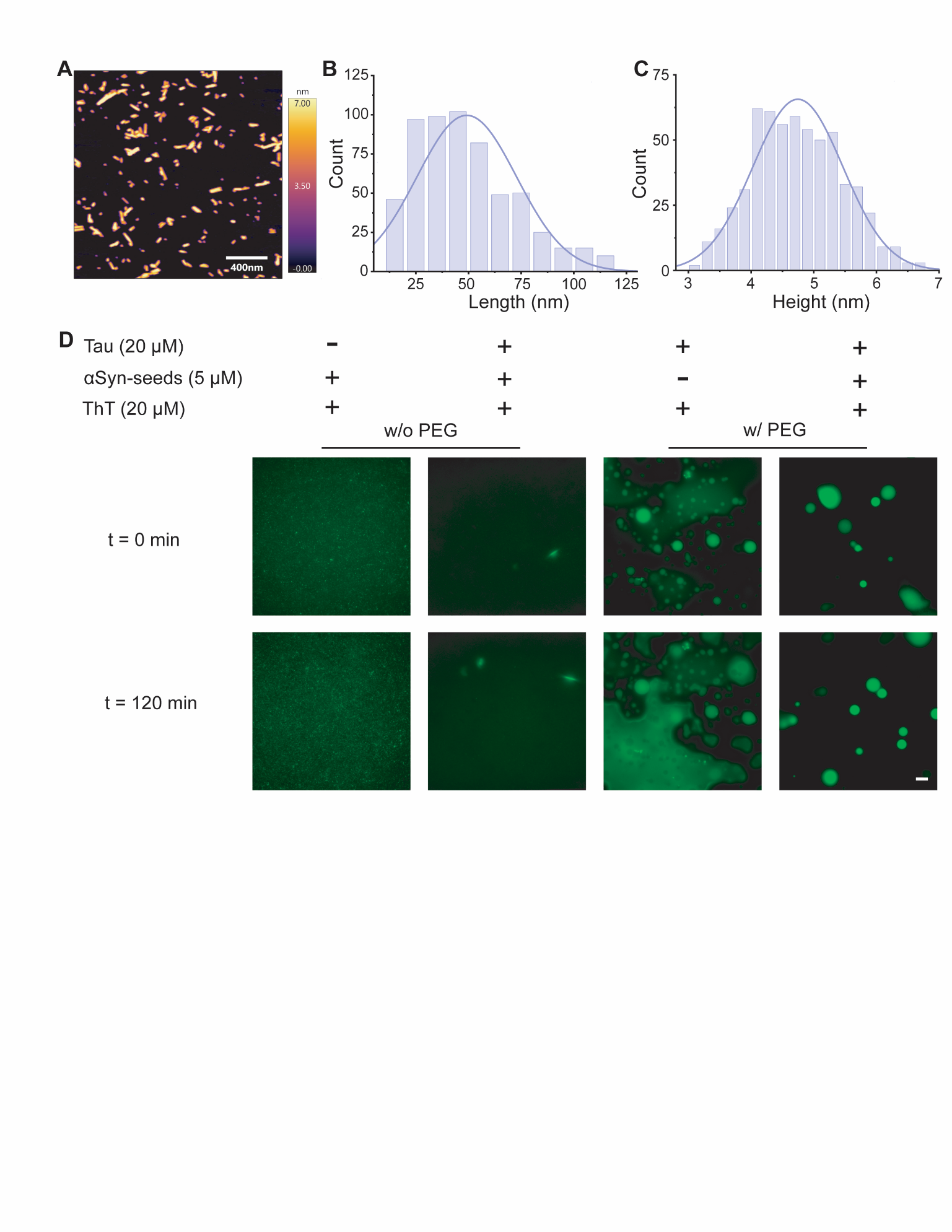
